# Dynamical Systems–Constrained Metabolic Modeling Enables Forecasting of Host-Microbiome Dynamics

**DOI:** 10.64898/2026.05.08.723020

**Authors:** Hayden Gallo, Vanni Bucci

## Abstract

Forecasting how microbiome-host ecosystems evolve through time simultaneously at the compositional and functional level remains a central challenge in biology. While dynamical systems models (DSMs) can infer and predict community composition from longitudinal abundance data, and constraint-based metabolic models (CBMMs) can estimate metabolic fluxes from genome-scale reconstructions, no existing framework unifies these approaches to generate mechanistically grounded, time-resolved forecasts of both microbial abundances and metabolite dynamics from ecological data alone. Here, we introduce the Dynamical Systems Constrained Metabolic Modeling (DySCoMeMo) framework, a new hybrid computational pipeline that integrates ecological DSMs with CBMMs to predict temporal dynamics of biomass and metabolites across microbial communities and hosts. DySCoMeMo leverages parameters inferred from application of DSMs to microbiome time series data to constrain metabolic modeling over time, thereby bridging ecological interaction networks with genome-scale metabolic modeling. DySCoMeMo is able to predict future community and metabolite dynamics in vitro with accuracy that is superior or on-par compared to that achieved with established methods that require actual microbial abundances and/or metabolites data for metabolite network inference or for estimating the per-microbe contribution to the extracellular metabolic pool. DySCoMeMo also generalizes to in vivo data as it is capable of accurately forecasting microbial and metabolite dynamics in response to dietary perturbations even when host metabolism is included. Finally, DySCoMeMo uniquely enables the identification of keystone species by quantifying their contributions to sustaining metabolic environments. Together, our work establishes a generalizable, mechanistically grounded framework for time-resolved forecasting of microbiome-host microbial and metabolic dynamics, bridging ecological interaction inference with genome-scale metabolism of communities.

## 1. Introduction

Intestinal microbiomes, the bacteria, their encoded genes and metabolic functions that live within the mammalian intestine, play a central role in shaping host physiology, immunity, and metabolism through metabolite-dependent dynamic interactions within the microbial community and with the host (1).

A central question in systems biology is whether we can quantitatively forecast how ecological interaction dynamics shape metabolic exchange within the microbiome and the host, and reciprocally, how metabolic constraints feed back to influence community structure over time.

Over the last decade, longitudinal dynamical systems models (DSMs) constrained by microbial abundance data, most prominently generalized Lotka–Volterra (gLV) formulations and their derivatives, have provided a powerful framework to capture species–species interactions and to reproduce temporal community dynamics (2, 3).

In these models, large systems of differential or difference equations are fit to temporal abundance data using approaches such as regularized regression (4–6), Bayesian inference (7, 8), and, more recently, neural ordinary differential equations (9), or physics-constrained neural networks (10).

Once parameterized, DSMs can be used to forecast microbial dynamics via numerical integration and solution of Initial Value Problem (IVP) (11) and to infer the ecological roles of specific taxa in shaping long-term ecological stability. Such interaction networks have revealed bacteria with growth-promoting or inhibitory effects on pathogens (6) and guided the design of bacterial consortia with immune-modulatory properties (5). These models have been also used successfully to assess species “keystones” (12) quantified as the impact that removal or introduction of a species or of a group of species has on community structure (7).

Despite their inherent advancements in moving microbiome analysis from purely descriptive to forecasting (2, 13), current DSMs approaches remain fundamentally agnostic to the metabolic functions encoded by each modeled microbial species. They capture who interacts with whom, but not how those interactions are mediated by metabolite exchange, nutrient competition, or host–microbiome co-metabolism.

Constraint-based metabolic modeling (CBMM) frameworks such as flux balance analysis (FBA) and flux variability analysis (FVA) have been shown to allow for prediction of metabolite production or consumption at steady state in microbial systems (14–18). Because of the underdetermined nature of the underlying system of linear equations, fluxes are solved through numerical optimization (i.e., linear programming) which requires the maximization of an objective function (14, 19). Traditionally, growth maximization has been shown to be accurate in reproducing metabolic dynamics in mono-culture experiments under unlimited nutrient conditions (20, 21), however this assumption may be inaccurate in multi-species microbial ecosystems where each microbial species’ effective growth rate depends on the presence and abundance of competitive microbes (22, 23). To address this, steady-state community metabolic modeling frameworks such as SteadyCom (24) and MICOM (25) have incorporated species abundance constraints to distribute growth rates across community members and predict community-level metabolic fluxes from metagenomic snapshots. However, these approaches operate at a single time point and cannot forecast longitudinal metabolic dynamics or account for ecological feedbacks between species over time.

Furthermore, CBMM still lacks the capacity to capture longitudinal dynamics and ecological feedbacks. Dynamic flux balance analysis (dFBA) has sought to extend CBMM into the temporal domain by coupling FBA with ordinary differential equations that track extracellular metabolite concentrations over time (22, 26). While dFBA can capture time-varying metabolite production and consumption, it typically assumes that microbial growth follows predefined kinetic parameters (e.g., Monod or Michaelis–Menten kinetics) and requires specification of exchange rates (usually*a priori* measured for aerobic model systems organisms, e.g., *Escherichia coli*) that are not available for complex host-associated microbiomes. As a result, dFBA has primarily been applied to single strains or very small consortia under well-characterized conditions (22, 26), rather than to the high-dimensional, host-associated microbial communities captured in longitudinal microbiome datasets. Recent dFBA-based simulators such as COMETS (27, 28) and *µ*BialSim (29) have sought to scale community metabolic modeling to larger consortia, yet still rely on pre-specified kinetic parameters and do not leverage ecological interaction inference from time-series data. Finally, no existing dFBA framework connects observed community dynamics inferred from longitudinal data (4–6) with mechanistic metabolic modeling predictions, leaving a fundamental gap between data-driven ecological inference and genome-scale metabolism.

To date, no unified framework enables mechanistically grounded, time-resolved forecasting of how ecological interaction dynamics shape metabolic exchange in complex host–microbiome ecosystems. Here we bridge ecological DSMs and CBMM to introduce the Dynamical Systems Constrained Metabolic Modeling (DySCoMeMo) framework. DySCoMeMo leverages ecological parameters inferred from DSM fitting to longitudinal microbial abundance data to impose time-resolved constraints, including time-dependent growth rates, equilibrium structure, and local stability, on multi-species genome-scale metabolic models. In this way, DySCoMeMo uses ecology to restrict the feasible metabolic space, allowing only those flux solutions that are consistent with the observed community dynamics while maintaining mechanistic biochemical realism.

By applying DySCoMeMo to recently published *in vitro* high-throughput co-culture datasets involving communities composed of one to twenty-five distinct taxa (30–32), we first demonstrate that it accurately predicts species abundances and metabolite concentrations for previously unseen conditions and without using the actual metabolite data for training.

We then apply DySCoMeMo to *in vivo* longitudinal datasets from animal experiments examining how diet shapes microbiome recovery following antibiotic perturbation (33). In this setting, DySCoMeMo successfully captures both the temporal trajectories of community composition and the overall distribution of key metabolites across the recovery period, highlighting its ability to integrate ecological dynamics with mechanistic metabolism in host-associated settings.

In contrast to ecological dynamics-based inference methods, DySCoMeMo allowed us in both the *in vitro* and *in vivo* cases to identify specific cross-feeding metabolites that mediate the observed ecological interactions which are consistent with known biological and experimental-based literature. Additionally, for the *in vivo* case DySCoMeMo allowed us to determine the relative contribution of specific bacteria and secreted metabolites to host metabolism over time.

Finally, DySCoMeMo can identify metabolic keystone species in complex microbial communities. By systematically evaluating predicted metabolite fluxes across all possible *n* − 1 microbial combinations (7), DySCoMeMo quantifies the contribution of individual taxa to sustaining key metabolic functions in the *in vitro* co-culture datasets described in (30– 32).

Altogether, DySCoMeMo becomes the first unified framework to link ecological dynamics to metabolic mechanisms within a microbiome–host ecosystem. By embedding DSM constraints into multi-species genome-scale metabolic models, it bridges the long-standing divide between data-driven ecological inference and mechanistic biochemical modeling. This integration enables DySCoMeMo not only to reproduce and forecast community and metabolite trajectories, but also to mechanistically resolve how individual species shape the metabolic environment within a microbiome.

## 2. Results

### 2.1. DySCoMeMo conceptual overview

A central conceptual insight underlying DySCoMeMo is that microbial communities do not operate in a single dynamical regime but instead move between periods of stability and transient behaviors (34–37). Classical theory predicts that complex ecosystems harbor a finite attractor landscape of stable steady states, with transient dynamics shaped by ecological interactions and perturbations (38). Yet no existing microbiome modeling framework has exploited this structure to constrain metabolic modeling predictions. DySCoMeMo does it explicitly: at each time point, it uses parameters inferred from DSM model fitting to time-series data (4, 7, 8, 10), to evaluate the local stability landscape of a microbial community (4, 7) and to determine whether this community resides near a stable equilibrium or traverses a non-stationary regime (4, 38). In stable regimes, DySCoMeMo enforces a SteadyCom-like metabolic solution (24) consistent with the equilibrium structure (See Methods); in transient regimes, it constrains metabolic growth using DSM-derived ecological rates, ensuring that flux solutions remain dynamically feasible. In this way, DySCoMeMo operationalizes the concept that metabolism in a microbial ecosystem must be interpreted through the lens of its ecological attractor structure. Below we briefly formalize the DySCoMeMo framework.

Given time-series abundance data for *n* distinct species in the community, ***X***(*t*) = [*x*_1_(*t*), *x*_2_(*t*), …, *x*_*n*_(*t*)], DySCoMeMo first fits a DSM of the form

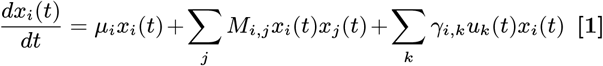

with *i* = 1, …, *n*, to infer the underlying ecological parameters **Θ** = [***µ***; ***M***; ***γ***], namely the intrinsic growth rates ***µ***, the pairwise interaction matrix ***M***, and the environmental perturbation coefficients ***γ*** associated with *p* measured external variables ***u***(*t*) = [*u*_1_(*t*), *u*_2_(*t*), …, *u*_*p*_(*t*)]. This inferential step is performed off-line and can be carried out using several established approaches (4, 6–8, 10, 30, 31). Importantly, DySCoMeMo is agnostic to the specific inference method used: any approach capable of accurately estimating ***µ*** and ***M*** from longitudinal abundance data can serve as the ecological foundation for the framework.

Once the ecological parameters **Θ** have been inferred, DySCoMeMo uses the parameterized system of equations to characterize the stability landscape of the community and to determine whether the ecosystem is residing near a stable steady state or traversing a transient, non-equilibrium regime (Fig. 1). Following standard gLV theory also applied to microbial community analysis (39), steady states correspond to those fixed points **x**∗ satisfying

**Fig. 1.**
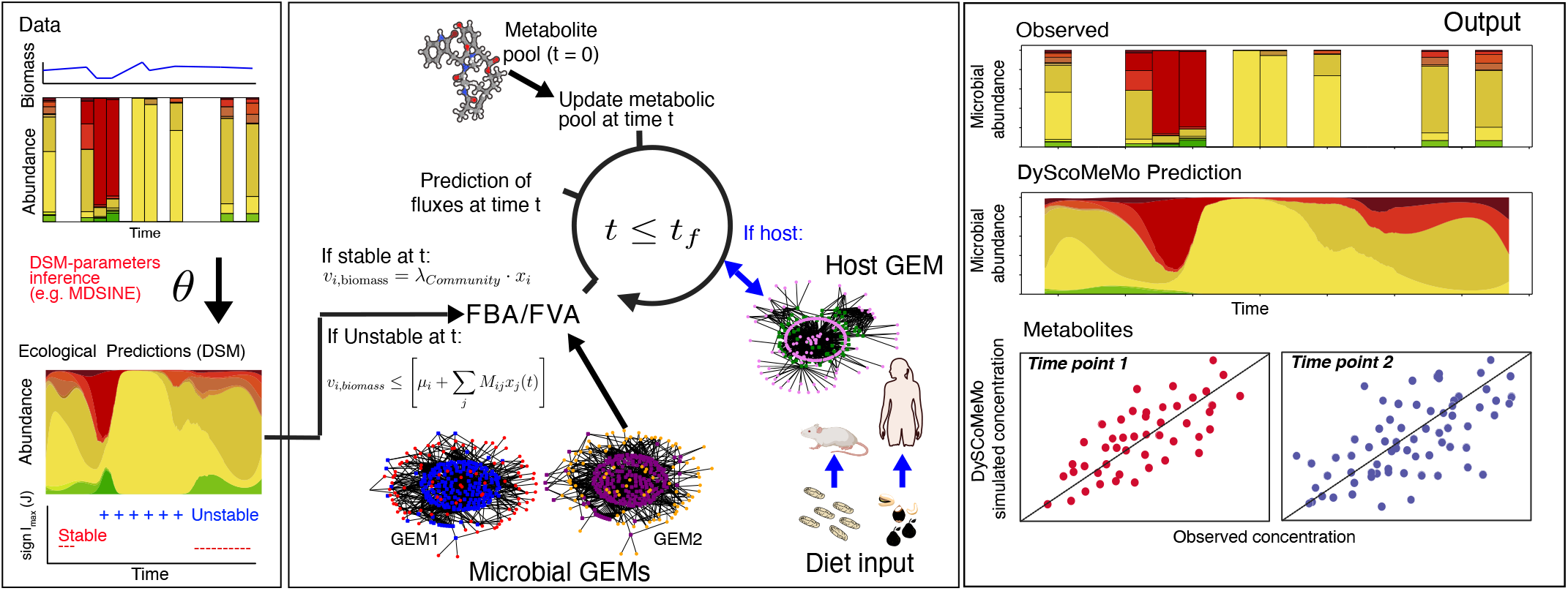
Overview of the DySCoMeMo framework. DySCoMeMo integrates ecological dynamical systems modeling with constraint-based metabolic modeling to generate time-resolved predictions of microbial abundances and metabolite concentrations. Longitudinal abundance data are first used to infer ecological parameters (growth rates and interaction matrix) using dynamical systems models (e.g., gLV via MDSINE/MDSINE2 (7, 8)). At each time point, DySCoMeMo evaluates local ecological stability: when the system is near a stable equilibrium, a SteadyCom-like formulation (24) is used to resolve community metabolic fluxes; when the system is in a transient or unstable regime, DSM-inferred per-capita growth rates are used to constrain species-specific biomass fluxes during FBA calculations. Predicted fluxes are then used to update the extracellular metabolite pool over time, optionally incorporating host genome-scale metabolic models and dietary inputs. This stability-aware switching enables DySCoMeMo to reconcile ecological dynamics with mechanistic metabolism, yielding accurate predictions of microbial abundances and metabolite trajectories across time.

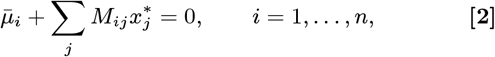

with 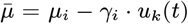, which are subject to the feasibility condition 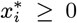. These fixed points represent candidate ecological equilibria toward which the system converges under appropriate environmental conditions.

To determine the dynamical regime at any given time, DySCoMeMo then performs a local stability analysis by evaluating the Jacobian of the DSM at each observed state **X**(*t*):

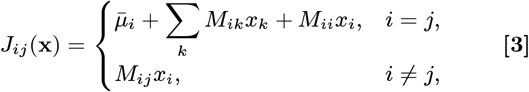

as in (4, 5, 7). The eigenvalues of *J* (**x**) characterize the local behavior: if all eigenvalues have negative real part, the system is locally attracted toward a stable equilibrium; if at least one eigenvalue has positive real part, the system is in a transient or unstable regime. This information is critical to DySCoMeMo, as it determines how metabolic fluxes are constrained at each time point. Specifically, DySCoMeMo uses these stability criteria in real time to classify the ecological state of the community. When the community is close to a stable fixed point (i.e., growth rates 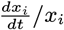 are near zero and the leading eigenvalue of *J* is negative), DySCoMeMo uses a metabolic mode which is in line with SteadyCom-like optimization. Here, the community composition is held constant, and the metabolic fluxes for each species are required to satisfy steady-state feasibility constraints. Here, the biomass production rate for each species is set to

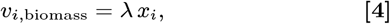

where *λ* is the common community growth rate which is equal to the highest achievable growth-rate by the least-performing microbial species. This constraint ensures that the predicted metabolic outputs reflect the long-term, stable behavior observed in the data under the scenario that all species would be growing at similar rate. This approach is justified in the original SteadyCom (24), which demonstrates that enforcing steady-state constraints accurately captures the metabolic outputs and species abundances observed in stable microbial communities. Conversely, when the system is in a non-stationary regime indicated by positive or near-zero eigenvalues, DySCoMeMo enters its transient regime, using the instantaneous ecological rates *µ*_*i*_ + ∑_*j*_*M*_*ij*_ *x*_*j*_ (*t*) to impose dynamic upper bounds on species-specific metabolic growth (Fig. 1)

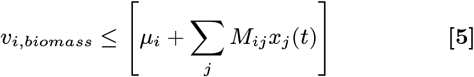

The dynamical systems model is necessary during transients: it identifies the current ecological regime and supplies the parameters needed to constrain metabolic fluxes away from equilibrium. By embedding these regime-aware constraints directly into flux balance optimization, DySCoMeMo ensures biological plausibility as community composition shifts, reproducing both steady-state and transient metabolic behaviors across diverse conditions

After pre-computing the ecological regime at each time point, DySCoMeMo runs a forward simulation integrating ecological dynamics with genome-scale metabolic modeling. Given an initial extracellular metabolite pool and dietary inputs, each species’ GEM is optimized subject to regime-appropriate constraints, SteadyCom-like at equilibrium, DSM-informed during transients, ensuring predicted fluxes remain consistent with both ecological dynamics and biochemical feasibility. A host GEM can optionally be included to capture host-microbiome metabolic exchange (Fig. 1)

The resulting fluxes update the extracellular metabolite pool at each time step, while microbial abundances are updated based on FBA-consistent growth rates, yielding the full system state. This procedure iterates forward, generating time-resolved predictions of both metabolite concentrations and community composition. All results presented here use FBA at each time step; DySCoMeMo is formulation-agnostic and can accommodate alternative constraint-based approaches such as parsimonious FBA (pFBA) (40) or FVA (41), though incorporating flux variability at each dynamically constrained time point would substantially increase computational cost and is left for future work.

Together, this regime-aware integration enables DySCoMeMo to reproduce both steady-state and transient metabolic behaviors, establishing a mechanistic link between community dynamics and metabolic function. By leveraging longitudinal ecological data alone, the framework generates time-resolved predictions of microbiome–host metabolic exchange and enables identification of species that govern ecosystem stability and metabolic output.

### 2.2. DySCoMeMo predicts microbial and metabolite dynamics in *in vitro* synthetic communities

We first evaluated the ability of DySCoMeMo to accurately predict microbial species abundances and metabolite dynamics over time, by applying it to two recently published *in vitro* co-culture datasets (30, 31). In these experiments, combinations of 25 common human gut anaerobes were grown under batch conditions in deep 96-well plates for 48 hours in defined media. Microbial abundances were quantified using 16S rRNA gene sequencing, while short-chain fatty acid (SCFA) concentrations were measured using targeted metabolomics. Measurements were collected either at the initial and final time points (0 and 48 hours; 761 community combinations, hereafter referred to as *in vitro Dataset 1*) or at multiple intermediate time points (0, 16, 32, and 48 hours; remaining combinations, hereafter *in vitro Dataset 2*). These datasets have served as the experimental foundation for several recent modeling studies. In the original work introducing *in vitro Dataset 1*, Clark et al. (30) jointly modeled bacterial abundances and metabolite concentrations using gLV dynamics for community composition and linear regression models to predict metabolite levels from species abundances, with the goal of guiding synthetic microbial community design. Subsequently, Baranwal et al. (31) demonstrated that recurrent neural network (RNN) models could further improve predictions of both species abundances and metabolite concentrations (for both *in vitro Dataset 1 and 2*) compared to the regression-based approaches used in (30). More recently, Quinn-Bohmann et al. (32) used endpoint bacterial abundances from *in vitro Dataset 1* to experimentally validate butyrate levels predicted using Microbial Community-scale Metabolic Models (MCMMs).

Crucially, in both Clark et al. (30) and Baranwal et al. (31), metabolite measurements were required for model training to infer species-specific contribution and to predict metabolite concentrations through time. Consequently, metabolite dynamics could not be predicted solely from microbial abundance data without prior metabolite observations.

On the other end, while in the MCMM-based analysis of (32), metabolite levels were predicted from bacterial abundances alone via GEMs, this was achieved statically (i.e., using the measured end-point bacterial abundances) rather than a true forecast of future states from previously-observed community compositions.

Compared to these approaches, DySCoMeMo relies exclusively on inferred parameters from longitudinal microbial abundance data and initial metabolite pool to forecast future microbial and metabolite levels through time that are consistent with the observed ecological dynamics. No metabolite observations are used to link species to metabolite levels, nor observed bacterial abundances.

We trained DySCoMeMo on the microbial abundance trajectories of *in vitro Dataset 2* and inferred the underlying gLV dynamics using Bayesian regression methods implemented in MDSINE2 (8) (Fig. 2A). Using the information from the MDSINE2-inferred gLV parameters as constraints on each individual species per-capita growth rate, we run DySCoMeMo’s metabolic modeling component to predict both species biomass and extracellular metabolites abundance from their experimentally-measured initial conditions (Fig. 2B,C). Because these experiments were conducted under batch conditions and do not exhibit steady-state behavior, we then operated DySCoMeMo exclusively in its DSM-constrained (transient) regime by imposing as maximum growth rate for every species the per-capita rate extracted from the gLV parameters, see eq. 5. DySCoMeMo was able to accurately rank species based on their abundance at each time point comparably to MDSINE2 (Fig. 2B) (Spearman’s *ρ >* 0.9; *p <* 1*e*^−9^). Additionally, DySCoMeMo was able to accurately rank the measured metabolites at each time point (Fig. 2C) (Spearman’s *ρ* ≥ 0.54; *p <* 1*e*^−^6).

**Fig. 2.**
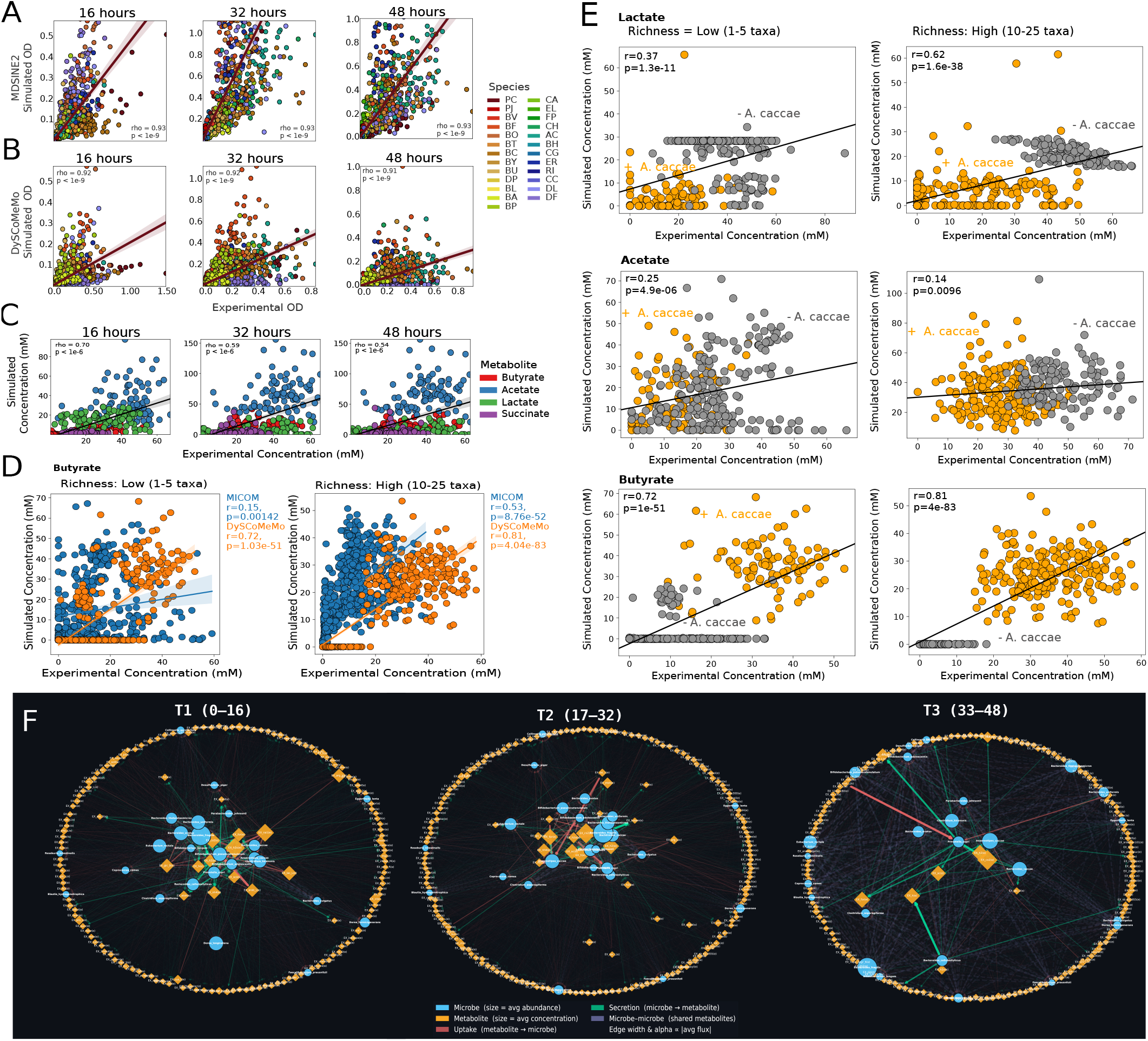
DySCoMeMo predicts microbial and metabolite dynamics in *in vitro* synthetic communities and reconstructs metabolite-mediated interaction networks. (A) Comparison between experimentally measured optical density (OD) and OD predicted from MDSINE2-inferred gLV dynamics at 16, 32, and 48 hours for communities in *in vitro Dataset 2*. Each point represents a species abundance in a given community; colors denote species identity. (B) Same as (A) but using DySCoMeMo, where metabolic fluxes are constrained by gLV-inferred ecological dynamics. DySCoMeMo reproduces the ranking of species abundances across time points with accuracy comparable to MDSINE2. (C) Predicted versus experimental metabolite concentrations (butyrate, acetate, lactate, succinate) at 16, 32, and 48 hours for *in vitro Dataset 2*. DySCoMeMo accurately ranks metabolite levels across communities using only microbial abundance–derived constraints. (D) Forecasting metabolite production for *in vitro Dataset 1*. DySCoMeMo was trained on *in vitro Dataset 2* and used to simulate community dynamics forward in time from the 0-hour initial conditions. Predicted butyrate concentrations at 48 hours are compared with experimental measurements for low-richness (1–5 taxa) and high-richness (10–25 taxa) communities. Performance is compared with the MCMM (MICOM) model reported in (32). (E) Predicted versus experimental concentrations of lactate, acetate, and butyrate at the 48-hour endpoint of *in vitro Dataset 1*. Points are colored according to the presence or absence of *A. caccae*, highlighting the role of *A. caccae* in determining extracellular metabolite levels. (F) Metabolite-mediated microbial interaction network inferred from DySCoMeMo flux predictions for *in vitro Dataset 2*. Directed edges represent metabolite exchange inferred from predicted secretion and uptake fluxes, linking producer and consumer species through shared metabolites. Because interactions are derived from genome-scale metabolic fluxes constrained by ecological dynamics, the inferred network captures mechanistic cross-feeding relationships across the full metabolic network rather than only measured metabolites.

Using the gLV parameters inferred from *in vitro Dataset 2*, we solved the Initial Value Problem (IVP) (11) and numerically integrated DySCoMeMo forward in time from the 0-hour initial conditions of *in vitro Dataset 1* (for both microbes and metabolites) to forecast microbial abundances and metabolite concentrations up to the 48-hour experimental endpoint.These predictions were evaluated against experimentally measured metabolite levels, enabling a direct assessment of DySCoMeMo’s ability to forecast metabolic outcome under previously unseen conditions (Fig. 2D,E). DySCoMeMo was able to accurately rank microbial tested combinations according to their resulting end-point butyrate concentration (Fig. 2D). DySCoMeMo achieved a predictive accuracy, evaluated as in the original paper, for both the low and high richness community scenarios for *in vitro Dataset 1*, see (32), which was higher compared to the MCMM model (MICOM) described in (32) (Fig. 2D). Low richness: (Pearson’s *r* = 0.72; *p* = 1.03*e*^−51^). High richness: (Pearson’s *r* = 0.81; *p* = 4.04*e*^−83^). Importantly, the MCMM-based analysis of (32) relies on the actual measured 48 hour endpoint abundance data to predict metabolites, while for DySCoMeMo this is a forecast from an IVP of a model trained on an independent dataset. DySCoMeMo was able to also accurately rank microbial compositions based on their outputted lactate and acetate concentrations at the 48 hour endpoint also with respect to lactate and acetate (Fig. 2E), a finding not reported in the original work by (32). Importantly, DySCoMeMo simulations clearly identify the presence or absence of the lactate-to-butyrate-converting bacterium *Anaerostipes caccae* as the main contributor to extracellular metabolite concentration in these experiments(Fig. 2E), a biological finding that clearly aligns with its known biological function (42), which was also described in (30).

Reconstructing metabolite-mediated microbial interaction networks has traditionally relied on correlation-based approaches applied to static microbiome abundance or metabolomic data (43). These methods identify statistical associations between species and metabolites that occur more frequently than expected by chance, but they do not provide mechanistic insight into the metabolic exchanges that drive microbial interactions. Temporal changes in metabolite-mediated interactions have also been explored using dFBA simulations(44), although these approaches typically rely on assumed kinetic parameters and are not calibrated against experimentally observed community dynamics. More recently, DSMs have been used to infer metabolite–microbe interaction networks directly from joint microbial and metabolite time-series data(10), however, such approaches are inherently limited to the subset of metabolites measured experimentally.

DySCoMeMo enables reconstruction of time-resolved metabolite-mediated microbial trophic interaction networks by identifying directed cross-feeding relationships from predicted secretion and uptake fluxes (Fig. 2F). Because these interactions are inferred from genome-scale metabolic models constrained by longitudinal ecological dynamics, the resulting networks extend beyond experimentally measured metabolites and provide a mechanistic view of community metabolic organization. In *in vitro Dataset 2*, DySCoMeMo recovered biologically plausible trophic structure. For instance, there was a lactate-mediated interaction linking *Roseburia intestinalis* and *Dorea formicigenerans* to *Anaerostipes caccae*, consistent with the established role of *A. caccae* as a lactate-utilizing, butyrate-producing bacterium (42) (Fig. 2F, SI Dataset S1, SI Dataset S2, Movie S1).

Likewise, multiple *Bacteroides* species, including *B. ovatus, B. thetaiotaomicron, B. uniformis*, and *B. vulgatus*, occupy upstream positions in the trophic web. These taxa are found by DySCoMeMo to be the primary consumers of a range of medium-enriched carbohydrate-derived metabolites consistent with these microbes’ recognized role as primary degraders of dietary carbohydrates in gut microbial communities (45). In addition to sugar uptake, several *Bacteroides* species were predicted to secrete intermediate metabolites including succinate, fructose, alanine, and proline, suggesting that they supply downstream substrates to other community members (Fig. 2F, SI Dataset S1, SI Dataset S2,Movie S1).

Finally, DySCoMeMo captured more specialized metabolic interactions beyond canonical SCFA cross-feeding, including sulfate uptake and hydrogen sulfide release by *Desulfovibrio piger* (Fig. 2F, SI Dataset S1, SI Dataset S2,Movie S1), a bacterium experimentally demonstrated to perform such metabolic function (30, 46), indicating that the inferred trophic structure encompasses both central carbon fermentation and sulfur-associated metabolism.

Altogether, these analyses establish that DySCoMeMo can accurately forecast microbial and metabolite dynamics from microbial abundance data alone by integrating ecological interaction inference with genome-scale metabolic modeling. Unlike existing statistical and constraint-based approaches, DySCoMeMo does not require metabolite measurements for training and produces time-resolved predictions that remain consistent with the underlying ecological dynamics of the community. In addition, by leveraging predicted secretion and uptake fluxes, DySCoMeMo enables mechanistic reconstruction of metabolite-mediated trophic interactions as they evolve through time, recovering interaction patterns consistent with known gut microbial biology.

### 2.3. DySCoMeMo accurately reconstructs longitudinal microbial and metabolite dynamics following antibiotic perturbation

To evaluate DySCoMeMo’s ability to recover host-associated microbiome and metabolome dynamics, we applied the framework to longitudinal *in vivo* data from a recent study by Kennedy et al. (33). In this work, the authors investigated microbiome resilience and functional recovery following broad-spectrum antibiotic perturbation in a mouse model. Longitudinal profiling revealed a structured successional recovery process characterized by dynamic shifts in taxonomic composition and metabolic function over time, enabling reconstruction of community reassembly and metabolite trajectories during restoration of ecological stability.

To apply DySCoMeMo to the *in vivo* dataset from Kennedy et al. (33), we first estimated the per-species total abundance in every sample by scaling the 16S rRNA-produced relative abundances by the Colony Forming Units (CFUs) measurements which quantify overall bacterial count, as in (4–7). The resulting per-species total abundance trajectories were used to infer gLV parameters via MDSINE2 (8), which yielded ecological interaction networks, growth rates and response to antibiotic perturbations as in (4–7).

DySCoMeMo used the inferred DSM parameters to then pre-compute the ecological regime (steady vs transient state) at every time point by solving equation eq. 3. This allowed us to determine what type of optimization would need to be enforced during the metabolic modeling calculations at each specific time point. Antibiotic contribution to the ecological and metabolic dynamics was implicitly accounted for by the ecological regime estimation because, as described above, 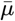 is the net growth rate which includes reduction during antibiotic treatment.

After resolving the ecological regime at each time point, the metabolic modeling module of DySCoMeMo was initialized at the pre-antibiotic steady state (time *t* = −3) using the observed community composition and corresponding initial metabolite profile (60 metabolites per diet; see Methods). Dietary information (e.g., composition of dietary sources and their concentration) was derived from (33), however because in our computation metabolic fluxes were estimated at an hourly resolution, daily dietary inputs were evenly distributed across each 24-hour period for simplicity.

In contrast to the *in vitro* application, the *in vivo* setting included the explicit inclusion of host metabolism. Accordingly, a GEM of *Mus musculus* was incorporated (47, 48), in which host metabolic fluxes were optimized at each time point and the resulting metabolite exchanges were used to update the shared metabolic pool before being passed to the microbial metabolic module. This coupling allowed DySCoMeMo to capture bidirectional host–microbiome metabolic interactions, and to generate time-resolved predictions of microbial and host-associated metabolite dynamics throughout the antibiotic perturbation and recovery period.

To evaluate model performance in reconstructing microbial population dynamics, we compared DySCoMeMo predictions against experimentally observed species abundances across longitudinal time points spanning pre-perturbation through recovery (Day −3 to Day 14).

As shown in Fig. 3A, predicted versus observed abundances exhibit strong concordance during early and intermediate recovery phases. Immediately following antibiotic treatment (Day 0-2), correlations remained high (Day 1: Spearman’s *ρ* = 0.793; Day 2: *ρ* = 0.864, *p <* 1*e*^−5^), indicating that the model successfully captures early successional restructuring of the community. During mid-to-late recovery (Days 4-7), predictive accuracy remained significant (Day 4: *ρ* = 0.776; Day 5: *ρ* = 0.700, *p <* 1*e*^−5^), although correlation strength gradually declined at later time points (Day 11: *ρ* = 0.43, p = 0.06; Day 14: *ρ* = 0.516, p = 0.02). This temporal trend suggests that while the model robustly captures early deterministic recovery dynamics driven by metabolic niche reoccupation, later stages likely incorporate host- or stochastic-driven processes not fully represented in the current mechanistic structure. Importantly, DySCoMeMo’s performance was comparable to that of MDSINE2 during early recovery (e.g., Day 1: MDSINE2 *ρ* = 0.924 vs DySCoMeMo *ρ* = 0.793), and remained statistically significant across most time points, supporting its ability to infer species-level dynamics using ecological-dynamics-informed constraints.

**Fig. 3.**
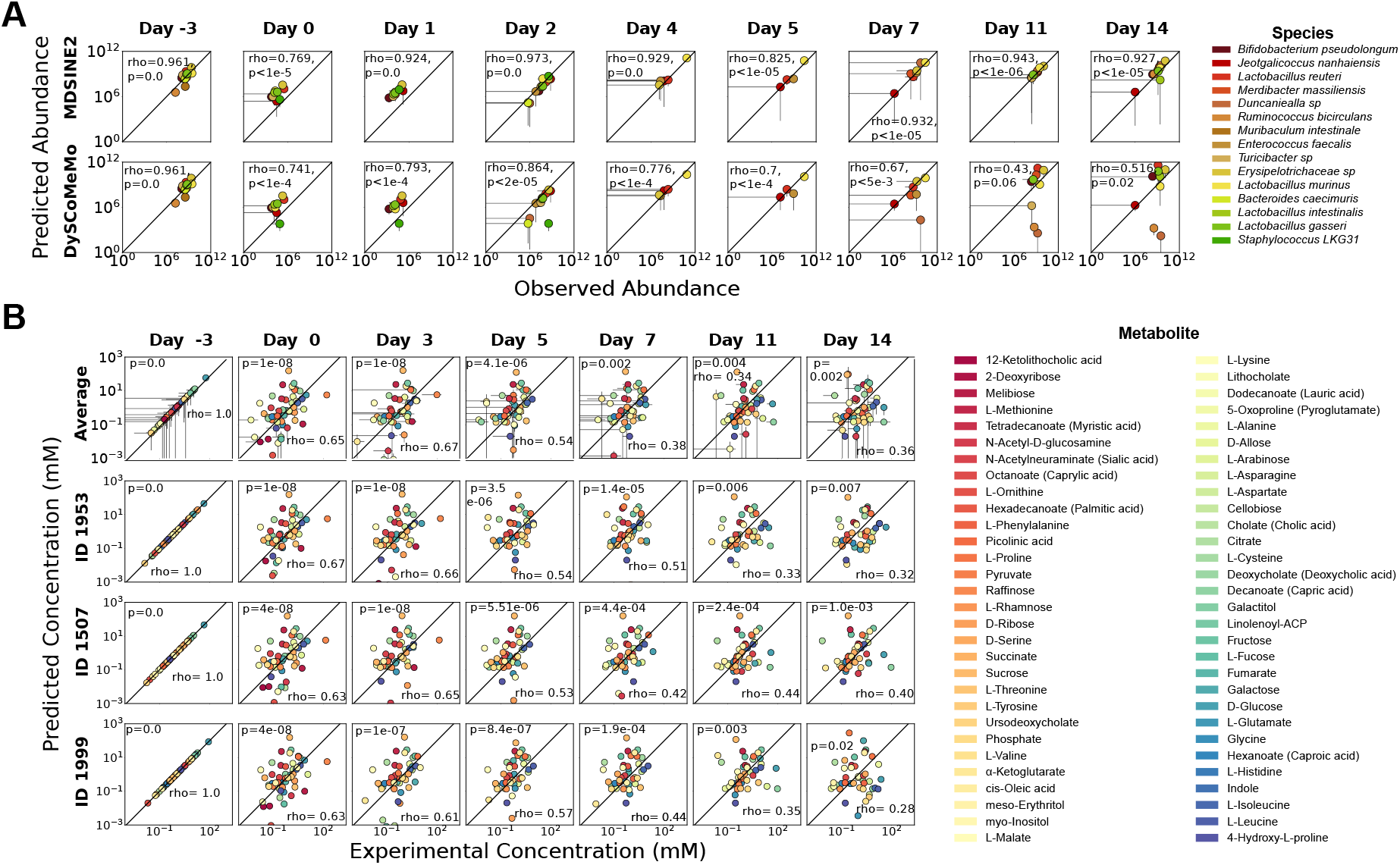
DySCoMeMo accurately reconstructs longitudinal microbial and metabolite dynamics following antibiotic perturbation. (A) Predicted versus observed species abundances across time points spanning pre-perturbation (Day −3) through recovery (Day 14). Each panel shows log-scale comparisons between model-predicted and experimentally measured abundances for individual species (color-coded). The diagonal line indicates perfect agreement. Spearman correlation coefficients (*ρ*) and associated p-values are reported for each time point. Top row: MDSINE2 predictions. Bottom row: DySCoMeMo predictions. DySCoMeMo achieves strong predictive performance during early and intermediate recovery phases, with gradual decline at later time points. (B) Predicted versus experimental metabolite concentrations (mM) across longitudinal time points. Top row: cohort-average predictions. Subsequent rows: individual subjects (IDs 1953, 1507, 1999). Each point represents a metabolite (color-coded), with error bars indicating experimental variability. Log-scaled axes and diagonal reference lines denote perfect agreement. Pearson correlations demonstrate strong predictive accuracy prior to perturbation and during early recovery, with reduced concordance at later stages.

We next evaluated the model’s ability to predict metabolite concentration trajectories across individuals and at the population-average level (Fig 3B). At the cohort-average level, model predictions showed significant correlations during early recovery (Day 0: *ρ* = 0.651; Day 3: *ρ* = 0.672, *p <* 1*e*^−8^). Correlation strength gradually declined during later recovery stages (Day 7-14: *ρ* = 0.34-0.38, *p <* 0.005), mirroring trends observed in species-level predictions. At the individual level (IDs 1953, 1507, 1999), DySCoMeMo captured inter-individual variability in metabolite trajectories with statistically significant correlations across most time points. For example, ID 1953 demonstrated strong early agreement (Day 0: *ρ* = 0.670; Day 3: *ρ* = 0.659, *p <* 1*e*^−8^), while ID 1507 maintained moderate predictive performance throughout recovery (Day 7: *ρ* = 0.418; Day 11: *ρ* = 0.437, *p <* 2*e*^−4^). Even in later stages, correlations remained significant in multiple individuals (e.g., ID 1999 Day 5: *ρ* = 0.573, *p* = 8.4*e*^−7^).

Collectively, these results demonstrate that DySCoMeMo accurately reconstructs metabolite dynamics during antibiotic-driven ecological disruption and early successional recovery, with declining but still significant performance during later, potentially host-modulated stabilization phases.

### 2.4. DySCoMeMo predicts functional keystoneness at both community-wide and metabolite-specific levels

Keystone species in microbial communities are organisms that exert a disproportionately large influence on community structure, stability, and function relative to their abundance (49–51). Predicting keystoneness *a priori* is therefore essential for understanding how microbial ecosystems respond to perturbations. Although recent work has broadened the keystone concept to include taxa or functions whose perturbation induces an outsized shift in community state, most current approaches still define keystoneness primarily at the compositional level, relying on abundance patterns, ecological network topology, or leave-one-out changes in community composition (7, 12, 35). By contrast, DySCoMeMo evaluates keystoneness in functional terms, quantifying how strongly species reshape community metabolic output through changes in metabolite production, consumption, and cross-feeding structure.

To illustrate this concept, we again analyzed the *in vitro* co-culture system reported by Clark et al. (30) (*in vitro* Dataset 1). For each observed community containing *n* = 3 to 25 species, we compared extracellular metabolite profiles with those measured in experimentally observed *n* − 1 subcommunities generated by removing one species at a time. For both the full community and each matched leave-one-out sub-community, DySCoMeMo was initialized using the corresponding experimental abundances, extracellular metabolite concentrations, and growth-medium composition, and simulated forward to the 48-hour endpoint. We then quantified the impact of removing each species as the root mean squared change in extracellular metabolite concentrations between each *n*-species community and its paired *n* − 1 sub-community.

At the species level, DySCoMeMo recovered broad trends in multimetabolite perturbation magnitude, with significant agreement between observed and simulated mean community-wide perturbations (Pearson’s *r* = 0.44, *p* = 2.9 *×* 1*e*^−2^; Fig. 4A). This analysis identified *A. caccae* (AC) as the strongest overall keystone species, showing the largest average effect on global community metabolic output across leave-one-out communities, whereas other species such as Dorea longicatena (DL), Dorea formicigenerans (DF), and Clostridium asparagiforme (CA) also exerted substantial but more context-dependent effects. Thus, DySCoMeMo captures functional keystoneness at the level of aggregate community metabolism.

**Fig. 4.**
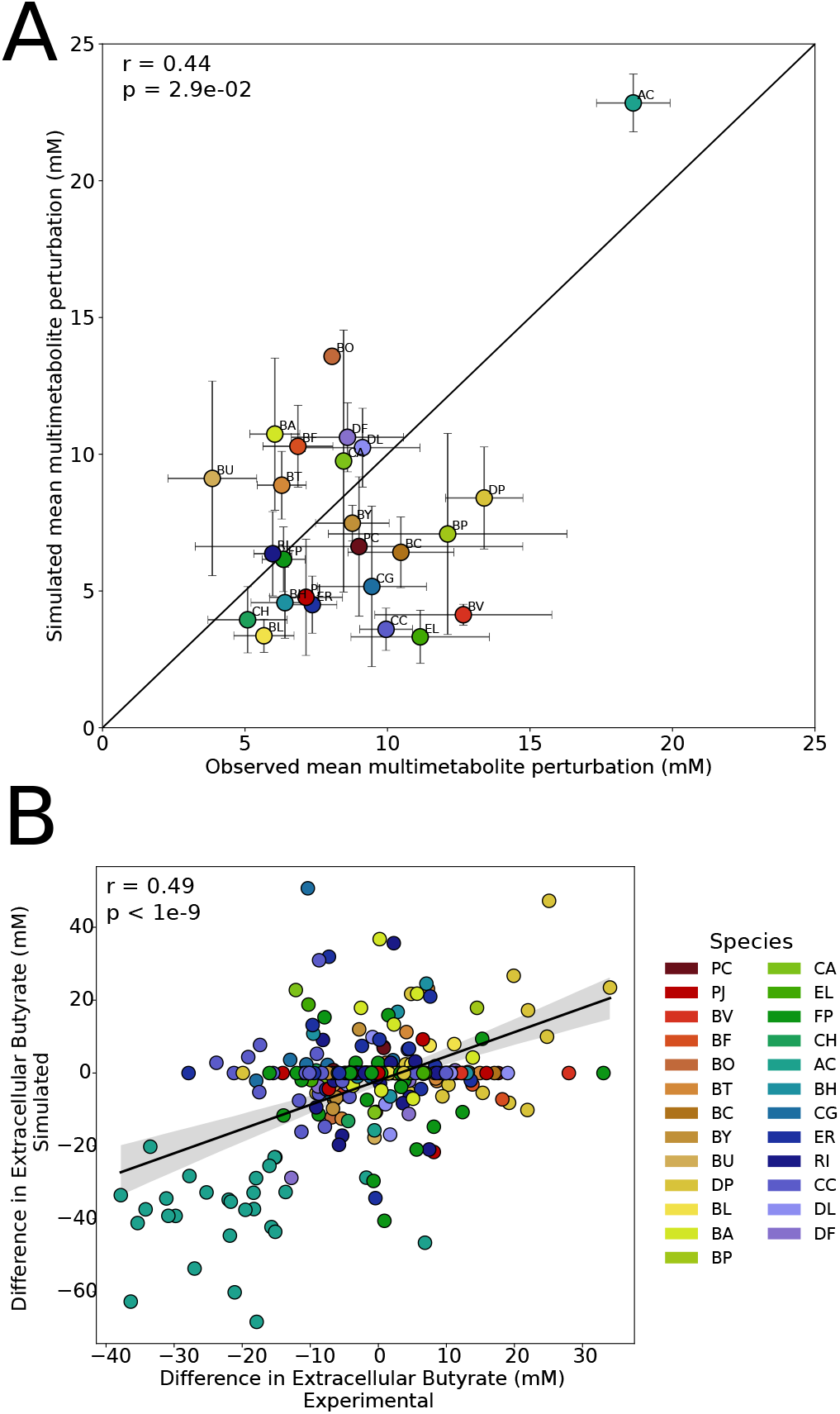
DySCoMeMo predicts functional keystoneness at both community-wide and metabolite-specific levels. **A**, Species-level comparison between experimentally observed and simulated mean multimetabolite perturbation following leave-one-out species removal across the *in vitro* Dataset 1 communities. For each species, perturbation was quantified as the root mean squared change in extracellular metabolite concentrations between each *n*-species community and the corresponding experimentally observed *n* − 1 sub-community obtained by removing that species. Points denote species-level means across all observed leave-one-out contexts, and error bars indicate variability across communities. The diagonal denotes perfect agreement between experiment and simulation. DySCoMeMo significantly recovered broad trends in functional keystoneness across metabolites, identifying *A. caccae* (AC) as the strongest overall keystone species. **B**, Comparison between experimentally observed and simulated leave-one-out effects on extracellular butyrate concentration across all communities. Each point corresponds to a single leave-one-out community. DySCoMeMo showed stronger agreement for butyrate-specific perturbations, supporting metabolite-specific keystoneness analysis and identifying *A. caccae* as the dominant butyrate keystone. DySCoMeMo also correctly identified *D. piger* (DP) as the strongest inhibitor of butyrate production.

We next asked whether DySCoMeMo could recover key-stoneness for a specific metabolic function. Focusing on butyrate, DySCoMeMo showed stronger agreement with experimentally observed leave-one-out effects (Pearson’s *r* = 0.49, *p <* 1*e*^−9^, *N* = 298; Fig. 4B), demonstrating that the model more accurately captures perturbations in this targeted metabolic output than in the aggregate multimetabolite phenotype. Under this metabolite-specific definition, *A. caccae* again emerged as the dominant keystone species for the control of community butyrate production. Further, *D. piger* was found to be the strongest reducer of butyrate production in these communities. These findings are consistent with the result in Clark et al. (30) that *D. piger* had the strongest negative impact on butyrate concentration whereas *A. caccae* had the most positive impact on the production of butyrate. Together, these results show that DySCoMeMo can quantify keystoneness both at the level of overall metabolic function and at the level of specific metabolic outputs.

## Discussion

A central challenge in microbiome systems biology is to move from descriptive associations to mechanistically-grounded forecasts of how microbial communities and their metabolic environments evolve through time (52). Ecological dynamical systems models can recover microbial interactions structure and predict community composition from longitudinal abundance data (4, 7, 8), whereas constraint-based metabolic models can estimate feasible metabolic states from genome-scale reconstructions (24, 25, 53), see also references in (54). However, as recently pointed out (55–57), these two modeling traditions have largely remained disconnected, leaving an important gap between ecological forecasting and mechanistic metabolic interpretation. In this work, we introduced DySCoMeMo to bridge this gap. By using parameters inferred from longitudinal ecological dynamics to constrain genome-scale metabolic modeling over time, DySCoMeMo provides a unified framework for forecasting both microbial abundances and extracellular metabolite dynamics while maintaining consistency with the observed ecological regime.

Our results show that this integration is useful across multiple levels of biological organization. In *in vitro* synthetic communities, DySCoMeMo accurately recovered time-resolved microbial and metabolite dynamics, and generalized to previously unseen community compositions without requiring metabolite measurements for model training. In doing so, it achieved performance that was superior or comparable to existing approaches that either require metabolite observations during training or rely on observed endpoint abundances rather than true forward forecasts. In addition, DySCoMeMo reconstructed metabolite-mediated trophic interactions that were consistent with known microbial biology, including lactic acid bacteria cross-feeding to *A. caccae*, upstream carbohydrate utilization by *Bacteroides* species, and sulfur-associated metabolism by *D. piger*. Extending the same framework to a host-associated setting, DySCoMeMo also recovered longitudinal microbial and metabolite dynamics during antibiotic perturbation and dietary recovery in vivo, supporting the idea that ecological constraints inferred from time-series data can be productively combined with mechanistic metabolic models even in more complex microbiome–host ecosystems.

Beyond prediction, an important conceptual advance of DySCoMeMo is that it makes it possible to quantify *functional* keystoneness. Most current keystone prediction methods in microbial ecology still operate primarily at the compositional level (7, 12, 35), relying on abundance patterns, ecological network topology, or simulated effects of taxon removal on community composition. Recent work has emphasized that keystoneness may also arise through metabolic functions and resource-mediated interactions (12), and has highlighted metabolic modeling as a promising route for keystone discovery. Our results operationalize that idea directly. By simulating matched leave-one-out communities, DySCoMeMo quantified the extent to which removal of an organism reshapes global metabolic output as well as specific metabolites. This analysis identified *A. caccae* as the strongest overall metabolic keystone in the synthetic gut community system examined here, while also showing that keystoneness depends on the functional phenotype under consideration: aggregate multimetabolite perturbation and butyrate-specific perturbation were related but not identical. In this sense, DySCoMeMo extends the keystone concept from “who changes community composition” to “who sustains community metabolic function,” which is likely to be more relevant for many microbiome-associated host phenotypes.

These results also help place DySCoMeMo within the broader modeling landscape. In contrast to purely ecological models, DySCoMeMo provides mechanistic hypotheses for *how* inferred interactions are mediated, by linking abundance trajectories to explicit secretion and uptake fluxes. In contrast to static community metabolic approaches, it enables time-resolved forecasting from initial conditions and can accommodate transient ecological regimes. And in contrast to standard dFBA implementations, it does not require the user to prescribe organism-specific kinetic parameters that are rarely available for host-associated microbiomes. Instead, the effective ecological constraints are learned from the longitudinal data themselves. We view this as a key strength of the framework: ecology is used to restrict the feasible metabolic space, while metabolism provides mechanistic interpretation of the ecological dynamics.

The present study still has several limitations. First, DySCoMeMo inherits uncertainty from both of its component models. Its performance depends on the quality of the inferred ecological parameters (58) and on the completeness and curation of the genome-scale metabolic reconstructions used for each organism (59). Errors in either layer can propagate into predicted metabolite trajectories and inferred trophic interactions. Second, the current implementation relies primarily on FBA with biomass-oriented objectives and does not yet propagate full flux uncertainty through time. Although the framework is compatible with alternative constraint-based formulations, exhaustive use of FVA or ensemble-based uncertainty quantification was beyond the scope of this study, as the associated computational cost scales with every simulation timestep (60). Third, host-associated simulations necessarily simplify many biological processes, including dietary exposure, host physiology, spatial compartmentalization, and regulatory effects that are not encoded in stoichiometric GEMs (61).

A natural extension will be to propagate uncertainty jointly across the ecological and metabolic layers, for example by coupling posterior samples from DSM inference to ensembles of metabolic models or flux solutions. It will also be important to incorporate richer host and environmental structure, including spatial compartmentalization, time-varying dietary inputs, and regulatory or thermodynamic constraints that refine feasible flux states. On the biological side, the functional keystoneness predictions generated by DySCoMeMo should be tested experimentally through targeted dropout, add-back, or metabolite rescue experiments, particularly in systems where multiple candidate keystones support overlapping functions. More broadly, integrating DySCoMeMo with intervention design frameworks could enable prospective prediction of how changes in diet, strain composition, or metabolite supplementation reshape both community structure and metabolic output.

Taken together, our results establish DySCoMeMo as a generalizable framework for linking ecological dynamics to metabolic mechanism in microbiome–host ecosystems. By unifying longitudinal dynamical systems inference with genome-scale metabolic modeling, DySCoMeMo makes it possible to forecast community and metabolite trajectories, reconstruct metabolite-mediated interactions, and quantify functional key-stoneness within a single mechanistic pipeline. We anticipate that this type of hybrid ecological–metabolic modeling will be an important step toward predictive, intervention-ready microbiome science.

## 3. Methods

### 3.1. Dynamical Systems Constrained Metabolic Modeling (DySCoMeMo)

DySCoMeMo is a hybrid computational framework that integrates ecological dynamical systems models (DSMs) with constraint-based metabolic models (CBMMs) to generate time-resolved predictions of microbial biomass and metabolite concentrations in microbial communities and host–microbiome ecosystems. The framework combines parameters inferred from longitudinal microbial abundance data with genome-scale metabolic models (GEMs) to constrain metabolic fluxes over time, ensuring consistency between ecological dynamics and biochemical feasibility.

At each time point, DySCoMeMo computes microbial growth rates using a generalized Lotka–Volterra (gLV) model inferred from experimental data and uses these rates to constrain flux balance analysis (FBA) of each species’ GEM. The resulting fluxes determine metabolite secretion and uptake rates, which are used to update the extracellular metabolite pool and microbial abundances at the next time step. This procedure is repeated iteratively to simulate community dynamics forward in time.

DySCoMeMo operates in two regimes depending on the ecological state of the system: a steady-state regime, in which community growth rates are balanced, and a transient regime, in which growth rates are constrained by the DSM. In this work, the transient regime was used for batch *in vitro* experiments, while both regimes can be used in host-associated systems.

### 3.2. Ecological dynamical systems inference

Microbial interaction parameters were inferred using the MDSINE2 framework (8), which fits generalized Lotka–Volterra (gLV) models to longitudinal microbial abundance data using Bayesian regression.

The gLV model employed by MDSINE2 is described by equation 1 in section 1.1 of the main text. Parameter inference in MDSINE2 is performed using a Bayesian hierarchical framework that accounts for measurement noise, biological variability, and uncertainty in model parameters. Posterior distributions over *µ*_*i*_ and *M*_*ij*_ and *γ*_*i,k*_ are obtained using Markov chain Monte Carlo (MCMC) sampling, allowing estimation of credible intervals for growth rates and interaction coefficients.

For each community, posterior samples of the gLV parameters were used to compute the instantaneous per-capita growth rate of each species at each time point as 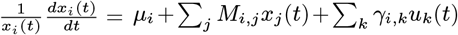. These growth rates were then used to constrain the metabolic optimization step in DySCoMeMo by imposing upper bounds on the biomass objective function of each species, ensuring that metabolic predictions remain consistent with the ecological dynamics inferred from experimental data.

### 3.3. Identification of ecological regime using fixed-point and Jacobian stability analysis

Once the ecological parameters (***µ, M***) were inferred from longitudinal abundance data, DySCoMeMo used the parametrized dynamical system to characterize the stability landscape of the microbial community and to determine whether the ecosystem was near a stable steady state or in a transient, non-equilibrium regime.

We consider the generalized Lotka–Volterra (gLV) model

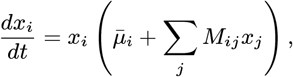

where *x*_*i*_ denotes the abundance of species *i, M*_*ij*_ represents pairwise ecological interactions, and

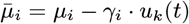

accounts for external perturbations or environmental inputs. To determine local stability conditions the procedure described in equations 2 and 3 in the section 1.1 of the main text was employed. DySCoMeMo uses this stability information to select the appropriate metabolic optimization regime at each time point.

#### 3.3.0.1. Steady-state regime

When the system is close to a stable fixed point, defined by negative real parts of all Jacobian eigenvalues and small per-capita growth rates, DySCoMeMo uses a steady-state community metabolic formulation consistent with the SteadyCom framework (24). This is further described in equation 4 and section 1.1 in the main text. This common community growth rate is the highest achievable growth-rate by the least-performing microbe, excluding species where *v*_*i,biomass*_ = 0. This formulation enforces steady-state feasibility while allowing metabolite exchange between species and environment.

#### 3.3.0.2. Transient regime

When the stability condition is not satisfied, indicating that the community is in a non-equilibrium regime, DySCoMeMo uses a dynamically constrained metabolic formulation.

In this case, the instantaneous per-capita growth rate predicted by the gLV model, see section 2.2, is used to impose an upper bound on the biomass reaction of each species: *v*_*i*,biomass_ ≤ *µ*_*i*_(*t*).

This constraint ensures that metabolic flux predictions remain consistent with the ecological dynamics inferred from the data and allows DySCoMeMo to simulate time-dependent changes in both species abundances and metabolite concentrations.

This regime-aware formulation allows DySCoMeMo to transition automatically between steady-state metabolic modeling and dynamically constrained metabolic modeling, enabling accurate simulation of both equilibrium and transient ecological conditions.

### 3.4. Genome-scale metabolic modeling

Each microbial species was represented by a genome-scale metabolic model (GEM). Metabolic fluxes were computed using flux balance analysis (FBA), which solves the optimization problem max_*v*_ *c*^*T*^ *v* subject to *Sv* = 0, *v*_*min*_ ≤ *v* ≤ *v*_*max*_, where *S* is the stoichiometric matrix, *v* is the vector of reaction fluxes, and *c* defines the biomass objective.

For each species, the biomass objective function was constrained by the per-capita growth rate predicted by the ecological model, 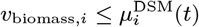.

This constraint ensures consistency between ecological dynamics and metabolic flux predictions.

All simulations in this study were performed using FBA at each time step. The DySCoMeMo framework is not restricted to FBA and can in principle accommodate alternative constraint-based formulations such as parsimonious flux balance analysis (pFBA) (40) or flux variability analysis (FVA) (41). However, performing FVA within a regime-aware, time-resolved community model would require solving a large number of additional optimization problems and was beyond the scope of this work.

### 3.5. Time-resolved simulation

DySCoMeMo simulations were performed by numerically integrating the coupled ecological– metabolic system forward in time.

At *t* = 0 the metabolite pool is initialized with experimental metabolite values. Then the forward simulation begins.

At each time step:

1. Per-capita growth rates were computed using the gLV model.
2. These rates were used to constrain the upperbound of the biomass objective function in each GEM.
3. FBA was solved for each organism.
4. Exchange fluxes were used to update extracellular metabolite concentrations.
5. Species abundances were updated according to the achieved FBA biomass rate and the species abundance at the beginning of the timestep.

Metabolites that have a concentration less than 10^−10^ mM are removed from the metabolite pool in order to avoid negative values and thus infeasible solutions. For *in vivo* simulations a decay term is employed to approximate washout of metabolites in the host intestine. For these *in vivo* simulations a decay term of 1 − (/48) was utilized.

The system was solved as an initial value problem (IVP) using numerical integration (11), starting from experimentally measured initial conditions.

### 3.6. In vitro co-culture datasets

DySCoMeMo was evaluated using previously published *in vitro* datasets (30, 31) consisting of synthetic microbial communities composed of 25 human gut anaerobes grown under batch conditions in defined media.

Microbial abundances were measured using 16S rRNA sequencing and metabolite concentrations were quantified using targeted metabolomics. When fitting the gLV MDSINE2 model to the microbial abundance data, the OD600 of each species was scaled by 10^8^ to improve the fit by MDSINE2.

For fitting, 25 random seeds with 10,000 MCMC samples each were used and then the outputs were concatenated and final inference was performed.

Dataset 1 contains measurements at 0 and 48 hours for 761 community combinations. Dataset 2 contains measurements at 0, 16, 32, and 48 hours for 95 community combinations.

DySCoMeMo was trained using bacterial abundance data from Dataset 2 to infer ecological parameters and evaluated by forecasting Dataset 1.

### 3.7. Forecasting of unseen communities

To evaluate predictive performance, gLV parameters inferred from Dataset 2 were used to simulate Dataset 1 by solving the IVP from experimentally measured initial conditions.

Predicted metabolite concentrations at 48 hours were compared with experimental measurements. Predictive accuracy was quantified using Pearson and Spearman correlation coefficients, following the evaluation procedure described in (32).

No metabolite measurements were used for model training.

### 3.8. Inference of metabolite-mediated interaction networks

Metabolite-mediated interaction networks were reconstructed from DySCoMeMo flux predictions.

At each time point, secretion and uptake fluxes were computed for all species. A directed interaction from species *i* to species *j* was inferred when a metabolite secreted by *i* was consumed by *j*.

Because fluxes were derived from genome-scale metabolic models constrained by ecological dynamics, the inferred networks represent mechanistic cross-feeding relationships across the full metabolic network rather than only experimentally measured metabolites.

### 3.9. Identification of metabolic keystone species

To quantify the contribution of individual species to community metabolic output, DySCoMeMo simulations were performed for all possible *n* − 1 species combinations (7).

For each community, metabolite production fluxes were computed and compared with those obtained in the full community.

A species was defined as a metabolic keystone if its removal caused a large reduction in the production of one or more key metabolites.

### 3.10. Application to in vivo host–microbiome datasets

DySCoMeMo was applied to longitudinal *in vivo* datasets in which microbial abundances and metabolite concentrations were measured during controlled perturbations.

Longitudinal microbial abundance data from 10 subjects were utilized for fitting the gLV MDSINE2 model. Each subject had 16S rRNA relative abundance data. These 10 subjects did not have CFU measurements but another cohort of subjects that underwent the same treatment schedule did. Therefore, utilizing this data and assuming a normal distribution, we randomly sampled to generate estimated CFU measures for our 10 subjects of interest.

Metabolomics data was not available for these subjects however, three of the ten subjects had cage pairs whom had metabolomics data available. Assuming well mixed conditions allowed for the use of these cage matched metabolomic and bacterial abundance data. These procedures are similar to the steps taken in Kennedy et al. (33).

Before fitting, the number of microbial species to be modeled was limited due to the feasibility of constructing genome scale metabolic models from scratch and to reduce matrix sparsity in fitting the gLV model. The filtering criteria applied was that a species must be present in at least one sample for all subjects. This filtering limited the number of species to model to 15.

Microbial interaction parameters were inferred using MDSINE2 from abundance time series. These parameters were used to constrain metabolic fluxes as described above. For fitting, 9 random seeds with 1,000 MCMC samples each were used and then the outputs were concatenated and a final inference was performed.

Once the gLV model was fit, we sampled the latent trajectory at an hourly resolution and smoothed using LOESS. To determine steady state or transient regimes in this data, we utilized 100 MCMC samples at daily resolution from the MDSINE2 produced latent trajectory. A *P* (*X >* 0.9) and relative abundance cutoff of 0.0001 were used as criterion for determining stable states.

Due to the fact that metabolic modeling works with abundances of grams of dry weight (gDW), when performing FBA total abundances were scaled accordingly. The scaling factor of 10^11^ for the *in vivo* dataset was utilized to allow for the balancing of microbial growth while allowing metabolites in the environmental pool to become sufficiently limited.

### 3.11. Host–microbiome metabolic modeling

Host metabolism was represented using a genome-scale metabolic model coupled to microbial GEMs through a shared extracellular compartment representing the intestinal lumen.

Exchange reactions allowed metabolites to be transferred between microbes, host, and environment.

Dietary inputs were incorporated by constraining nutrient uptake reactions according to the experimental diet.

Perturbations such as antibiotic treatment were modeled by modifying ecological parameters, which in turn constrained allowable biomass fluxes in the metabolic optimization.

Minimal metabolites were added explicitly and only for host consumption to allow for minimal growth in the dietary conditions.

Further, host biomass was limited to an upper bounded growth rate of 0.5*hr*^−1^.

### 3.12. Model evaluation

Predicted microbial abundances and metabolite concentrations were compared with experimental measurements at each time point.

Accuracy was quantified using Pearson and Spearman correlation coefficients across species and metabolites.

All predictions were generated without using metabolite measurements for training.

### 3.13. Genome scale metabolic model utilization and reconstruction

For both *in vitro* datasets the same metabolic models were used. Here the AGORA2 database of genome scale models were used at the panGenus level. This is consistent with the methods described by Quinn-Bohmann et al (32).

In the case of the *in vivo* dataset, genome scale metabolic models were reconstructed using KBase (62). For reasons similar to what was described previously (33), we did not use AGORA2 reconstructions, specifically due to the inability to gapfill AGORA2 models on the dietary conditions of this study. We performed reconstructions on the species level, annotated utilizing RAST (63), and created draft GEMs via the MS2-Build Prokaryotic Metabolic models with OMEGGA online application in KBase (62). Following gapfilling procedures used in (33), models were gapfilled on a media composition to eliminate auxotrophies and also on a glucose minimal media. Models were also gapfilled on the dietary conditions that included the minimal media metabolites found in base KBase medias. Literature and bacterial growth data was surveyed to determine if each individual species could grow on fructose, sucrose and/or raffinose which were all dietary components in this study. If a species was found to grow on one of these carbon sources, the reconstruction was further gapfilled in a minimal media with the aforementioned carbon source. Models were then exported from KBase.

### 3.14. Media reconstruction

We performed different media constructions for the *in vitro* and *in vivo* datasets. For the *in vitro* dataset we translated the compounds for the defined medium (DM38) that was utilized by Clark et al. (30) to perform the synthetic community experiments. To ensure that all panGenus AGORA models could grow in the media, we performed a relaxed FBA (64) where we ensured each model could achieve a minimal growth rate of 1*h*^−1^. The minimal number of needed metabolites to achieve this for all models were added to the media formulation at a concentration of 0.1 mM. For dietary conditions in the *in vivo* modeling, we used experimental metabolomics data from Kennedy et al. (33) that characterized the concentration of the 60 metabolites of interest in the RC diet. The dietary data was treated as mmol/g concentrations and KBase minimal media at concentrations of 1.0 mM were further added to the diet reconstruction. Dietary components were scaled by a factor of 5, to reflect an estimated daily intake of 5g and then these values were evenly spread across 24 hours. For all media construction, water was assumed to be non-limiting.

### 3.15. Software and implementation

All simulations were implemented in Python using COBRApy for constraint-based modeling, MDSINE2 for ecological inference, and custom code for the DySCoMeMo framework.

Optimization problems were solved using the Gurobi solver. Numerical integration was performed using SciPy.

All code used in this study will be made available upon publication.

### 3.16. Data, Materials, and Software Availability

Data and code will be deposited to https://www.github.com/haydengallo/glvdfba. All other data is available in the manuscript and/or the supplementary information. Previously published data was also used for this work (30, 31).

## ACKNOWLEDGMENTS

Funding for this project was provided by the Gates Foundation INV-046284, by the National Institute of Allergy and Infectious Disease U01AI1729875, and by the National Institute of Aging R01AG075283 to Vanni Bucci. Hayden Gallo acknowledges support by the NIH T32 AI095213-15.

